# SPARKLE: evidence-constrained correction of local RNA leakage in high-resolution spatial transcriptomics

**DOI:** 10.64898/2026.08.12.744394

**Authors:** Shuai Wang, Bolin Zhu, Shanyou Li, Xiaoyu Wei

**Author notes:** These authors contributed equally to this work (Shuai Wang, Bolin Zhu).

## Abstract

High-resolution sequencing-based spatial transcriptomics, including Stereo-seq and Visium HD, aggregates dense capture units into cell-resolved expression matrices. During tissue processing and permeabilization, RNA released from source cells can spread to neighbouring capture locations, reducing cell-type specificity and biasing downstream analyses. Here we developed SPARKLE Ambient RNA Kernel-based Leakage Estimator), a cell-level correction method that uses capture locations outside cell-segmentation masks as within-sample spatial evidence of leakage. SPARKLE sparse spatial kernels to out-of-mask observations to estimate a sample-level propagation scale and gene-specific leakage coefficients. It corrects only genes supported by out-of-mask goodness of fit and uses expression-dependent conservative shrinkage to protect highly expressing source cells. In ten simulated scenarios, SPARKLE achieved the highest cell-wise concordance in eight and the lowest RMSE in nine. In axolotl brain, mouse brain and human ovarian cancer, SPARKLE removed ectopic marker signal from neighbouring cells while retaining source-cell expression, improved agreement with independent single-cell and single-nucleus references, and recovered an inferred fibroblast-to-tumor *COL1A2-SDC4* communication route that was obscured by ectopic *COL1A2* expression. Conclusions remained stable across plausible spatial scales and background-bin sizes. Runtime scaled with tissue-window area and was further accelerated on GPU. SPARKLE is therefore a reference-free, fast and scalable method for correcting local RNA leakage from evidence contained within each improving the reliability of cell-type localization, tissue-compartment identification and cell-cell communication inference.

## Introduction

Spatial transcriptomics measures transcript abundance together with tissue position, allowing cell states to be analysed in situ [1]. High-resolution sequencing-based platforms such as Stereo-seq and Visium HD use dense arrays to capture transcripts at cellular or subcellular scales [2–5]. With image- or transcript-density-based segmentation, minimal capture units can then be aggregated into cell-resolved expression matrices for cell-type localization, spatial-neighbourhood analysis, tissue-boundary detection and cell-cell communication inference [5–9].

Tissue processing and permeabilization can release cellular contents and allow RNA to migrate locally, so extracellular transcripts may be captured by neighbouring barcodes [10]. The resulting contamination varies across space: genes that are abundant in a local source are more likely to appear at nearby positions, and the signal often decreases with distance. Leakage also varies by gene because abundance, stability, release propensity and capture efficiency differ, making a single contamination fraction unsuitable [10–12].

Local RNA leakage can reduce expression contrast between cell populations, reorder differential-expression results and spread signals beyond the tissue compartments in which they originate. Cell annotation, spatial-variation analysis and cell-cell communication inference often depend on low-abundance markers, so even modest contamination may change qualitative conclusions [7–9, 13]. Spatial proximity can also reflect genuine tissue transitions or biological interactions. Correction therefore needs direct evidence for the component being removed and should avoid subtracting high-confidence endogenous expression.

Existing methods address related forms of contamination but make different assumptions. SpotClean models spot swapping between adjacent capture locations and uses spatial coordinates in the contamination process [11]. It was designed for multicellular-resolution spot-based data and corrects capture spots rather than leakage among segmented cells in high-resolution spatial transcriptomics. SPLIT targets transcript spillover in imaging-based platforms such as Xenium by combining a single-cell reference with cell-type deconvolution [12], and SPIDER denoises spatial transcriptomic data through embedding regularization with single-cell supervision [14]; both methods require the single-cell reference to cover the relevant cell types and states. Ambient-RNA methods developed for single-cell data, such as SoupX, use empty droplets to estimate a background profile but generally do not model contaminant sources as a function of spatial distance in tissue [15].

High-resolution spatial transcriptomic data also contain capture locations outside cell-segmentation masks. These locations may include tissue debris, segmentation errors or nonspecific capture, but their coordinates can also provide direct within-sample evidence of RNA leakage from nearby cells [16, 17]. This makes it possible to estimate spatial decay and gene-specific leakage from out-of-mask observations without an external single-cell reference. The remaining challenge is to use that evidence without excessive subtraction during cell-level correction.

We developed SPARKLE (Spatial Ambient RNA Kernel-based Leakage Estimator) for high-resolution sequencing-based spatial transcriptomic data with minimal-capture-unit coordinates and cell-segmentation labels. SPARKLE keeps each segmented cell intact and aggregates capture locations outside cell masks into spatial bins. A truncated spatial kernel links source cells to these out-of-mask bins, and their observed counts are used to estimate a sample-level spatial decay scale λ and gene-specific leakage coefficients αg. SPARKLE then predicts local leakage on a cell-cell graph and uses expression-dependent conservative shrinkage to limit subtraction from highly expressing cells. This design targets locally supported background while retaining genuine spatial signal.

## Materials and methods

### Overview of SPARKLE

SPARKLE corrects local RNA leakage only when observations outside segmented cell masks support that leakage (Fig. 1). The method takes raw gene-by-capture-location counts, two-dimensional coordinates and cell-segmentation labels. Segmented cells are treated as potential leakage sources and as correction targets, whereas out-of-mask locations are used for parameter estimation.

**Figure 1.**
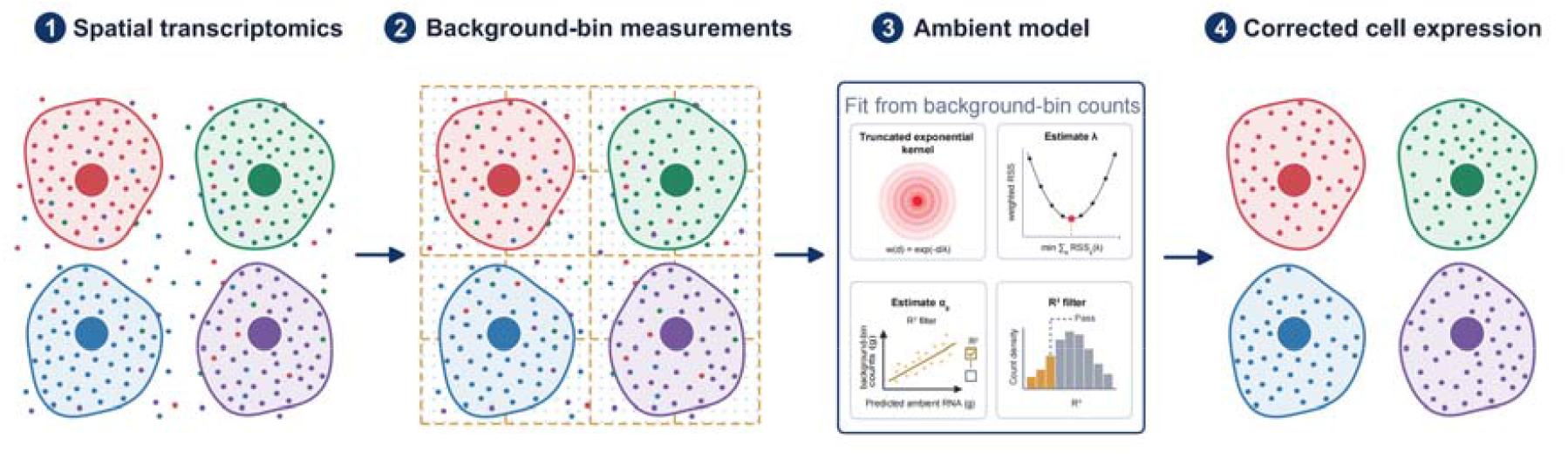
SPARKLE workflow for estimating and correcting local RNA leakage from out-of-mask spatial observations. Input high-resolution spatial transcriptomic data contain minimal-capture-unit coordinates and cell-segmentation labels. Intact cells are potential leakage sources and correction targets, whereas out-of-mask capture units are aggregated into background bins. The background-bin-to-cell graph estimates the shared propagation scale λ and gene-specific leakage coefficient α_g_; per-gene goodness-of-fit gating defines the correction set. A zero-diagonal cell-cell graph predicts local leakage, and expression-dependent conservative shrinkage protects highly expressing source cells to produce a non-negative corrected matrix.

The workflow has six stages:

1. obtain cell-level expression, effective area and centroids;
2. aggregate out-of-mask capture units on a regular grid to obtain background-bin counts, effective areas and centroids;
3. construct a sparse background-bin-to-cell distance graph and select a sample-level spatial decay scale from informative genes;
4. estimate gene-specific leakage coefficients at the selected scale and use a goodness-of-fit threshold to select genes for correction;
5. construct a zero-diagonal cell-cell graph and predict local leakage into each target cell from neighbouring cells; and
6. apply expression-dependent conservative shrinkage to limit subtraction from highly expressing cells, then truncate corrected values at zero.

### SPARKLE model

#### Inputs, outputs and statistical objective

Let a sample contain G genes and N coordinate-resolved minimal capture units (for example, DNBs or spots). The raw counts form a gene-by-capture-unit matrix denoted *X*, whose entry x_gi_ is the UMI count of gene g at capture unit i. The capture-unit coordinate is r_i_ = (r_il_,r_i2_), and its segmentation label is *ℓ*_i_: *ℓ*_i_ =c ≥ 0 indicates that capture unit i belongs to cell c, whereas *ℓ*_i_ −1 indicates a location outside all cell masks.

The model returns the corrected gene-by-cell matrix 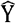, together with the estimated spatial scale 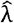, gene-specific coefficients 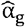, per-gene goodness of fit 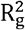 and the set of corrected genes. Because kernel predictions and regression coefficients generally yield non-integer leakage estimates, 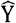 is a continuous non-negative matrix.

The direct statistical objective of SPARKLE is to fit the following relationship on out-of-mask background bins:

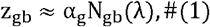

where z_gb_ is the observed total count of gene g in background bin b, and N_gb (_λ) is the spatial-kernel-weighted prediction from nearby segmented cells. SPARKLE does not include an additional global constant for background unrelated to observed neighbouring cells and therefore corrects only the locally explainable component.

Notation used in the SPARKLE framework is summarized in Table 1.

**Table 1.** Summary of notations used in the SPARKLE framework.

| Symbol | Description |
| --- | --- |
| $G, N$ | Number of genes and number of coordinate-resolved minimal capture units (for example, DNBs or spots), respectively |
| $X$ | Raw gene-by-capture-unit UMI count matrix |
| $x_{gi}$ | UMI count of gene $g$ at capture unit $i$ |
| $g, i, c, b$ | Indices of genes, capture units, cells and background bins, respectively |
| $r_i = (r_{i1}, r_{i2})$ | Spatial coordinate of capture unit $i$ |
| $\ell_i$ | Segmentation label of capture unit $i$: equal to $c$ ($\geq 0$) when the unit belongs to cell $c$, and $-1$ when it lies outside all cell masks |
| $\hat{Y}$ | Corrected gene-by-cell expression matrix (continuous and non-negative) |
| $z_{gb}, N_{gb}(\lambda)$ | Observed total count of gene $g$ in background bin $b$ and its spatial-kernel-weighted prediction at candidate scale $\lambda$, respectively |
| $\mathcal{I}_c$ | Set of capture units covered by cell $c$ |
| $a_c, \mu_c$ | Effective-area proxy (number of covered capture units) and centroid of cell $c$, respectively |
| $y_{gc}, \hat{y}_{gc}$ | Raw and corrected counts of gene $g$ in cell $c$, respectively |
| $s_{gc}$ | Source intensity of gene $g$ in cell $c$ per unit effective area |
| $h$ | Width of the out-of-mask background-bin grid |
| $\mathcal{E}_b$ | Set of observed out-of-mask capture units falling in background bin $b$ |
| $m_b, q_b$ | Effective-area proxy and centroid of background bin $b$, respectively |
| $K_\lambda(d)$ | Truncated exponential distance kernel; $d$ is the Euclidean distance between two coordinate pairs |
| $\lambda, R$ | Spatial decay scale of the leakage kernel and maximum neighbourhood radius of the spatial graphs, respectively |
| $\mathcal{G}$ | Candidate gene set for model fitting and correction |
| $\mathcal{G}_\lambda$ | Subset of candidate genes used for shared-scale estimation |
| $n_{\text{high}}, n_\lambda$ | Numbers of top-ranked genes retained in the candidate set and in the shared-scale subset, respectively |
| $\Lambda$ | Candidate set of spatial scales |
| $\alpha_g$ | Gene-specific non-negative leakage coefficient |
| $\text{RSS}(\lambda)$ | Joint area-weighted residual sum of squares at candidate scale $\lambda$ |
| $R_g^2, \tau_{R^2}$ | Per-gene goodness of fit and its threshold for entering correction (default 0.01), respectively |
| $\mathcal{G}_{\text{corr}}$ | Set of genes entering correction |
| $L_{gc}, A_{gc}$ | Kernel-weighted source intensity received by target cell $c$ from other cells and the final leakage estimate for gene $g$ in cell $c$, respectively |
| $p_{90,g}, f_{gc}$ | 90th percentile of the positive source intensities of gene $g$ across cells and the expression-dependent conservative shrinkage weight, respectively |
| $\mathbf{Y}^{\text{true}}$ | Contamination-free ground-truth cell-level expression matrix in simulations |
| $\alpha_0, U_g$ | Base leakage coefficient and the Uniform(0, 1) random variable used to generate true leakage coefficients in simulations, respectively |
| $\alpha_g^{\text{true}}, \lambda^{\text{true}}$ | True gene-specific leakage coefficient and true spatial decay scale in simulations, respectively |

#### Cell and out-of-mask observations

Denote by *J*_c_ the set of capture units covered by cell c. The effective-area proxy, centroid and total gene count of the cell are, respectively,

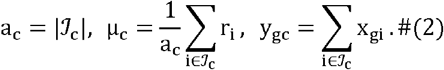

a_c_ is the number of minimal capture units covered by the cell mask. Within a platform, if every capture unit has a constant physical area, a_c_ is proportional to physical area. The source intensity per unit effective area for gene g in cell c is

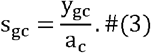

Out-of-mask capture units are aggregated on a regular grid of width ε_b_. Let be the set of observed out-of-mask capture units falling in background bin b; then

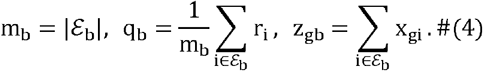

m_b_ is the effective-area proxy of the background bin, and q_b_ is the centroid of its observed locations. Grid cells containing no out-of-mask capture units are excluded. Edge bins and bins intersected by cell masks may therefore have different values of m_b_.

#### Sparse spatial graphs and distance kernel

By default, SPARKLE uses the truncated exponential kernel

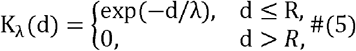

where λ > 0 controls the decay rate and R > 0 is the maximum neighbourhood radius. The exponential kernel provides a low-parameter, monotonic approximation to local decay.

SPARKLE constructs a background-bin-to-cell cross-graph and a cell-cell graph. The first connects every pair (b,c) satisfying d(q_b_,µ_c_) ≤ R and is used to estimate λ and α_g_; the second connects cell pairs c ≠ c ′satisfying d(µ_c_,µ_c_ ′)≤ R and is used for final correction. Graph topology and edge distances are constructed using KD-tree radius queries.

#### Selection of informative genes

Low-abundance genes usually contain many zero counts outside cell masks and provide little information for stable estimation of the spatial scale. Genes are therefore ranked by their mean count per unit effective area outside cell masks, computed as the total out-of-mask count of the gene divided by the total effective area of the observed out-of-mask bins.

The candidate set for model fitting and correction is denoted *G*. If n_high_ is specified, the top n_high_ ranked genes are used; otherwise, all genes are retained. The subset used for shared-scale estimation is *G*_λ_ ⊆ *G* and comprises the top n_λ_ genes in the candidate set.

#### Estimation of the shared spatial decay scale

For a candidate scale λ ∈ Λ, the unscaled source prediction for gene g in background bin b is

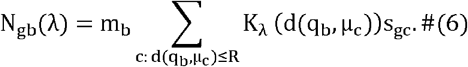

s_gc_ represents expression per unit effective area in a source cell, the spatial kernel represents distance-dependent contribution and m_b_ converts the prediction per unit area to the total-count scale of the background bin.

At fixed λ, a non-negative zero-intercept coefficient is fitted for every g ∈ G_λ_ by minimizing the area-weighted residual sum of squares between the observed and predicted background-bin counts, giving

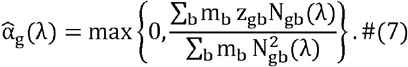

The coefficient 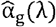 is re-estimated for every candidate scale, and the joint residual sum of squares is

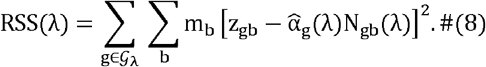

The selected scale 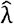 is the candidate value with the smallest joint residual sum of squares.

The shared λ constrains the shape of spatial decay using multiple informative genes, whereas differences in leakage magnitude between genes are represented by α_g_.

#### Gene-specific leakage coefficients and goodness-of-fit gating

After fixing 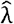, the coefficient 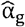 is re-estimated for every g ∈ *G* using the same expression as above.

The per-gene goodness of fit 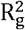 is the coefficient of determination of this area-weighted fit, computed from the area-weighted residual and total sums of squares of the background-bin counts. Candidate genes whose goodness of fit 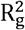 reaches a threshold 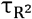 enter correction; they are denoted collectively by *G*_corr_, and the default threshold is 0.01.

#### Leakage prediction

For g ∈*G*_corr_, the kernel-weighted source intensity received by target cell c from other cells is

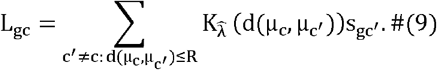

#### Expression-dependent conservative shrinkage and final correction

Spatially clustered cells of the same type can genuinely co-express a gene at high levels. To reduce overcorrection in this setting, standard SPARKLE uses expression-dependent conservative shrinkage. Let p_90,g_ denote the 90th percentile of the positive source intensities of the gene across cells, and define the shrinkage weight

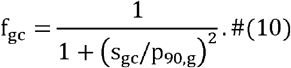

If the source intensity of gene g is zero in every cell, define f_gc_ = 1. The weight f_gc_ approaches 1 in low-expression target cells, whereas f_gc_ is smaller in high-expression target cells, thereby reducing subtraction from putative source cells.

The leakage estimate scales the received source intensity with the leakage coefficient and the target-cell effective area, which maps the intensity per unit effective area back to the total-count scale of the target cell. The final leakage estimate and corrected count are

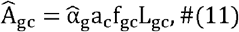

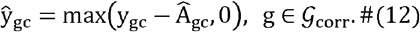

For g ∉ G_corr_, set ŷ _gc_ = y_gc_.

### Benchmark datasets

#### Simulated data

Simulations used a regular 500 × 500 DNB grid with 250,000 capture units and a DNB pitch of 0.5 µm, corresponding to an approximately 250 × 250 µm field of view. Each dataset contained 500 genes, including 80 highly expressed genes, with log-spaced expression gradients across genes so that most genes were expressed at very low levels. The target cell count ranged from 400 to 1,400, the mean cell radius was 5 µm and its coefficient of variation was 0.2. Each scenario contained 6 to 8 cell types with imbalanced proportions and strictly cell-type-specific marker genes. A 20% UMI dropout was applied to mimic capture loss, and the correction ground truth was defined as the observed post-dropout clean expression.

A contamination-free cell-level expression matrix Y^true^ was first generated and allocated to intracellular DNBs in proportion to cell area. Local leakage was injected only for highly expressed genes. Gene-specific true coefficients were generated as

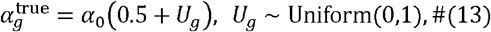

and Poisson leakage counts were injected into other cells and background DNBs using an exponential distance kernel with true scale λ ^true^. All simulations used random seed 42.

The ten predefined scenarios varied cell density, background fraction, spatial decay scale, leakage magnitude, marker fraction, number of cell types or spatial domain structure (Table 2):

**Table 2.** Parameter configurations for the ten simulated benchmarking scenarios.

| Scenario | Target cells | Background fraction | True $\lambda$ ($\mu\text{m}$) | Base leakage coefficient | Cell types | Setting |
| --- | --- | --- | --- | --- | --- | --- |
| S1 | 600 | 0.40 | 50 | 0.01 | 6 | Low density |
| S2 | 900 | 0.25 | 50 | 0.01 | 6 | Medium density |
| S3 | 1400 | 0.10 | 50 | 0.01 | 6 | High density |
| S4 | 900 | 0.25 | 20 | 0.01 | 6 | Short-range leakage |
| S5 | 900 | 0.25 | 500 | 0.01 | 6 | Long-range leakage |
| S6 | 900 | 0.25 | 50 | 0.01 | 6 | Spatially clustered cell types |
| S7 | 900 | 0.25 | 50 | 0.10 | 6 | Strong leakage |
| S8 | 400 | 0.60 | 50 | 0.01 | 6 | Very sparse tissue |
| S9 | 900 | 0.25 | 50 | 0.01 | 6 | Marker fraction increased to 50% |
| S10 | 900 | 0.25 | 50 | 0.01 | 8 | More cell types |

#### Real datasets

The real-data benchmarks covered the Stereo-seq and Visium HD high-resolution sequencing-based platforms [2, 5]. The axolotl brain, mouse-brain T304 and human ovarian-cancer data were obtained from published atlases and public data resources [18–23]. The principal analysis windows and available references are summarized below (Table 3).

**Table 3.**
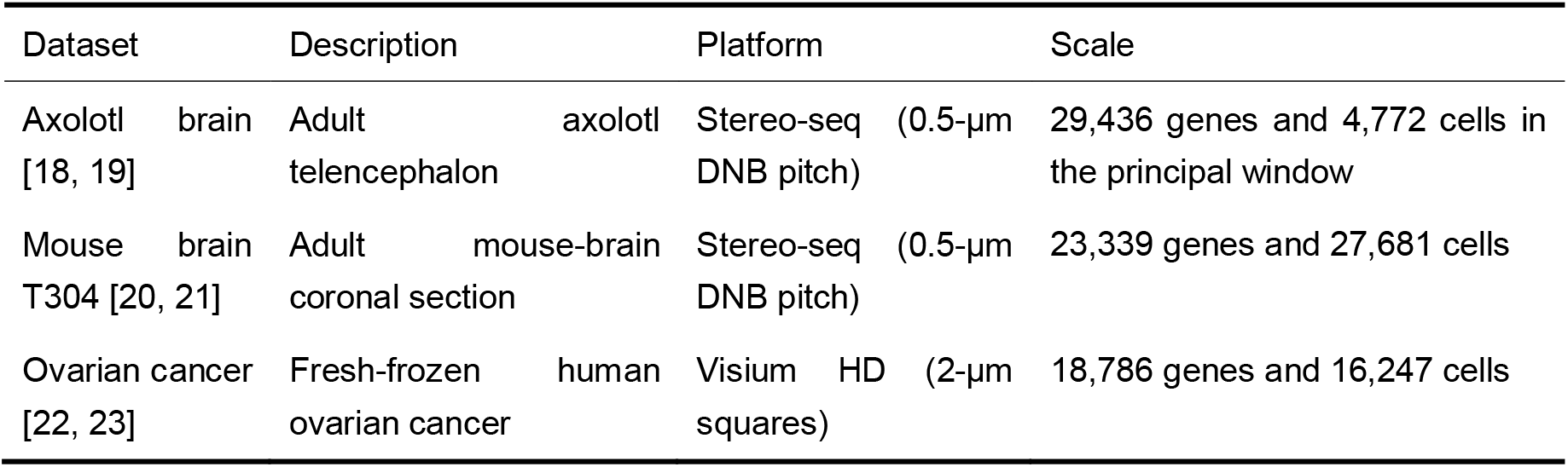
Overview of real-data benchmark datasets.

For the axolotl dataset, DNBs inside cell masks were aggregated by cell label, and CIDs absent from the label mapping were treated as out-of-mask observations. In the mouse-brain dataset, valid cells were defined by the T304 cell annotations; CIDs labelled 0 or not matching a valid cell ID were treated as out-of-mask locations. For the Visium HD ovarian-cancer dataset, the cell-segmentation mask was read directly, and 2-µm squares not covered by a mask were used as out-of-mask locations.

### Benchmark methods

#### SoupX

SoupX was developed to remove ambient RNA from droplet-based single-cell RNA sequencing [15]. For the spatial adaptation, the minimal-capture-unit matrix was supplied as the unfiltered table of droplets (tod), and capture units sharing a cell label were summed to form the cell-level table of counts (toc). Out-of-mask capture units were retained in the tod to estimate the background expression profile. SoupX was run using the official R package (v1.6.2): k-means clustering with a fixed random seed, contamination estimation with autoEstCont and count correction with adjustCounts. For simulations, tfidfMin = 0.05, soupQuantile = 0.5, quickMarkers FDR = 0.1 and the cluster number matched the true number of cell types; for real datasets, tfidfMin = 0.1, soupQuantile = 0.7, quickMarkers FDR = 0.01, with 6 clusters for axolotl and 30 clusters for mouse brain and ovarian cancer. forceAccept = TRUE was used throughout; the contamination estimate was capped at 0.8 for simulations and axolotl and at 0.99 for mouse brain and ovarian cancer.

#### DecontX

DecontX was run at cell level without spatial coordinates or out-of-mask observations [24]. Minimal capture units were first aggregated by cell label into a raw gene-by-cell count matrix. DecontX was run using the official R package celda (v1.26.0). Cluster labels were obtained by k-means clustering with a fixed random seed on log1p-transformed counts, with the number of clusters set to max(2, number of cells/20). Estimation used at most 200 iterations; cells with zero library size were excluded during estimation and restored afterwards.

#### SpotClean

SpotClean was run with the official Bioconductor R package (v1.12.0) [11]. To adapt it to cell-resolved data, each intact segmented cell was treated as one tissue spot; its count vector was the raw sum across covered capture units, and its coordinate was the arithmetic centroid of those units. Out-of-mask capture units were aggregated on a fixed grid to form background spots, with raw summed counts and centroids of the observed out-of-mask locations. SpotClean was run with kernel = “gaussian”, maxit = 30 and tol = 1.0.

### Benchmark metrics

Simulated data were evaluated against the contamination-free cell-level ground truth using two scale-robust metrics: the root-mean-square error (RMSE) on per-cell library-normalised log1p(CP10K) counts, and the Pearson R2, across genes, between corrected and true expression profiles for every cell. Results across scenarios are reported as mean ± SD.

For the axolotl *SST* analysis, annotated sstINs formed the source-positive group. A non-sstIN cell was assigned to the neighbouring group when its centroid was within the prespecified radius of any sstIN; all remaining non-sstIN cells formed the other group. We report the mean count in each group.

For mouse brain and ovarian cancer, cell-type expression profiles were constructed by averaging log-normalized expression across cells within each cell population. We calculated pairwise Pearson correlations between cell-type mean expression profiles, as well as correlations over shared genes between spatial profiles and the corresponding snRNA-seq or scFFPE single-cell reference profiles. Cell populations in mouse brain and ovarian cancer were assigned with RCTD [7] using the corresponding snRNA-seq or scFFPE single-cell reference.

For the ovarian-cancer marker analysis, counts were normalized to CP10K (Counts Per 10,000) and log1p transformed per cell before tumor-versus-stroma log2 fold changes were calculated for curated tumor and stromal markers. Spatial and single-cell CellChat analyses used their respective applicable modes [8].

### Computational performance and statistical reporting

CPU/GPU resource benchmarks used eight progressively larger mouse-brain windows. We recorded end-to-end runtime, the peak increment in host memory, and peak allocated and reserved GPU memory. Sensitivity analyses either fixed candidate λ values or changed background-bin side length while keeping other inputs and random seeds constant. Summary values across the ten scenarios are reported as mean ± SD; cell- or population-level distributions are shown as individual observations and box plots. All GPU analyses were run on a single NVIDIA GeForce RTX 4090 with CUDA 12.4, cuDNN 9.1.0, and PyTorch 2.6.0.

## Results

### SPARKLE improves cell-level expression recovery in simulations

We evaluated correction performance in ten simulated scenarios that varied cell density, background fraction, spatial decay scale, leakage magnitude, marker-gene fraction, number of cell types and spatial domain structure. For each cell, agreement between corrected and true expression profiles was measured as Pearson R2 across all genes. SPARKLE had the highest mean cell-wise R2 in 8 of 10 scenarios and averaged 0.892 ± 0.159 (mean ± SD) across all scenarios, compared with SoupX (0.884 ± 0.110), SpotClean (0.846 ± 0.146), DecontX (0.832 ± 0.211) and RAW (0.750 ± 0.195) (Fig. 2 a).

**Figure 2.**
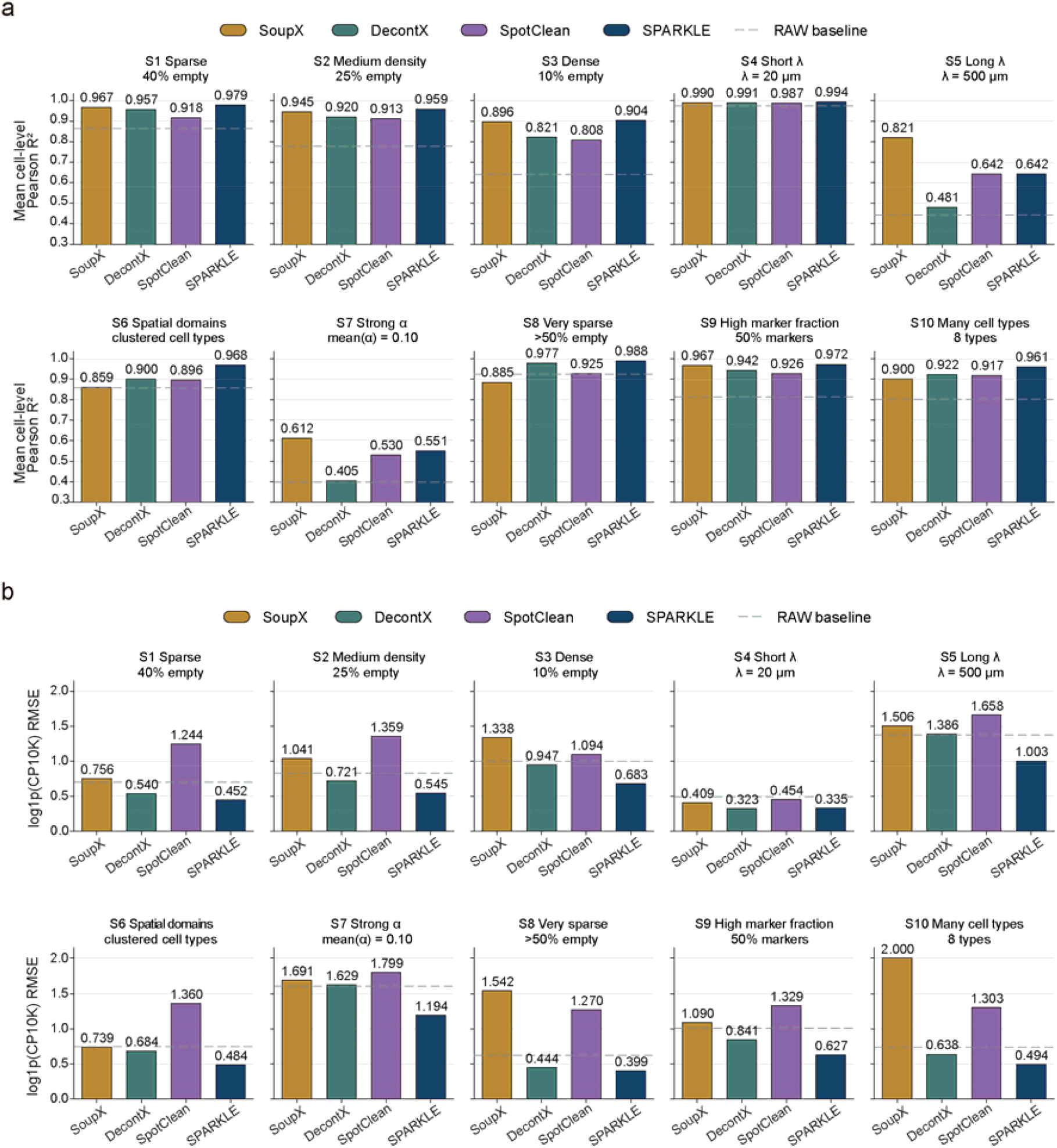
Cell-wise expression concordance across ten simulations. (a) Mean per-cell Pearson R^2^ with ground-truth expression across genes. (b) RMSE between corrected and ground-truth log1p(CP10K) expression. Higher R^2^ and lower RMSE indicate better recovery.

On the log1p(CP10K) RMSE scale, SPARKLE achieved the lowest error in 9 of 10 scenarios and reduced RMSE by 32.8% on average relative to RAW, compared with DecontX (14.1%). SoupX (−39.1%) and SpotClean (−47.1%) increased RMSE relative to RAW, likely due to overestimated ambient fractions. SpotClean inferred bleeding rates of 0.27-1.00 and redistributed subtracted counts; SoupX ρ estimates spanned 0.01-0.80, with several scenarios reaching the preset upper bound and requiring forceAccept (Fig. 2 b).

### SPARKLE reduces ectopic *SST* signal while retaining source expression

*SST*-expressing inhibitory neurons (sstINs) are sparse in the cortical region of the axolotl brain, so *SST* provides a useful marker for examining local RNA leakage (Fig. 3 a and b) [18]. The analysis window contained 206 sstINs, 505 non-sstIN cells neighbouring an sstIN and 4,061 other non-sstIN cells. In RAW data, mean *SST* counts were 73.0, 9.7 and 2.1 in these three groups, respectively, consistent with a signal that decreased with distance from sstINs (Fig. 3 c). After SPARKLE correction, the means were 69.9, 4.0 and 0.8. SPARKLE retained approximately 95.8% of the mean source-cell signal, reduced the mean signal in neighbouring cells by approximately 59.2%, and increased the sstIN-to-neighbour mean ratio from 7.53-fold to 17.69-fold (Fig. 3 c). SoupX and DecontX produced smaller reductions in neighbouring cells. SpotClean increased the mean sstIN count to 209.2 while retaining relatively high background signal in neighbouring and other cells.

**Figure 3.**
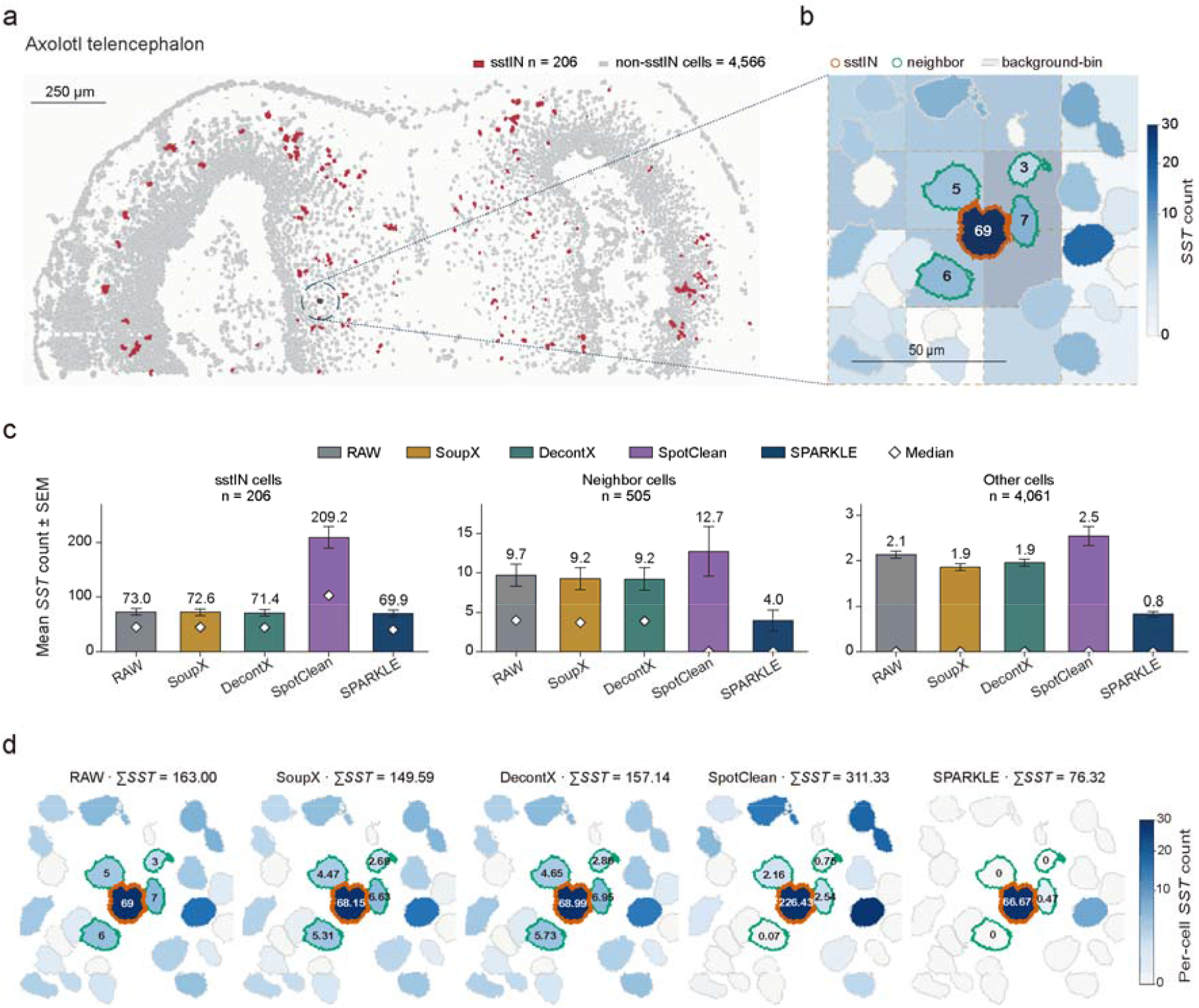
Retention of *SST* source-cell signal and removal of neighbouring ectopic expression in axolotl telencephalon. (a) Spatial distribution of sstINs in the analysis window. (b) *SST* expression in RAW data. (c) Mean *SST* counts ± SEM (bars and error bars) and medians (diamonds) in sstINs, neighbouring non-sstINs and other non-sstINs. (d) Per-cell *SST* counts in a representative sstIN neighbourhood across RAW, SoupX, DecontX, SpotClean and SPARKLE data.

At the level of a single neighbourhood, the example sstIN contained 69 *SST* counts in RAW data, whereas several adjacent non-sstIN cells contained 3 to 7 counts. After SPARKLE correction, the source cell retained 66.67 counts and signals in neighbouring and surrounding low-expression cells were markedly reduced (Fig. 3 d). The correction was concentrated on spatially dispersed *SST* rather than applied as a uniform reduction across cells.

### SPARKLE improves cell-type separation and reference concordance in mouse brain

In mouse brain, we compared separation among spatial cell types with concordance to an independent snRNA-seq reference [20]. Mean pairwise Pearson correlation between cell-type expression profiles decreased from 0.881 in RAW data to 0.833 after SPARKLE correction, consistent with less shared expression across cell types; among the other methods, DecontX produced a slightly larger decrease (0.825) whereas SoupX left the correlation essentially unchanged (0.881) (Fig. 4 a). At the same time, mean correlation between spatial cell-type pseudobulks and the snRNA-seq reference increased from 0.388 to 0.414, the largest improvement among the tested methods (Fig. 4 b).

**Figure 4.**
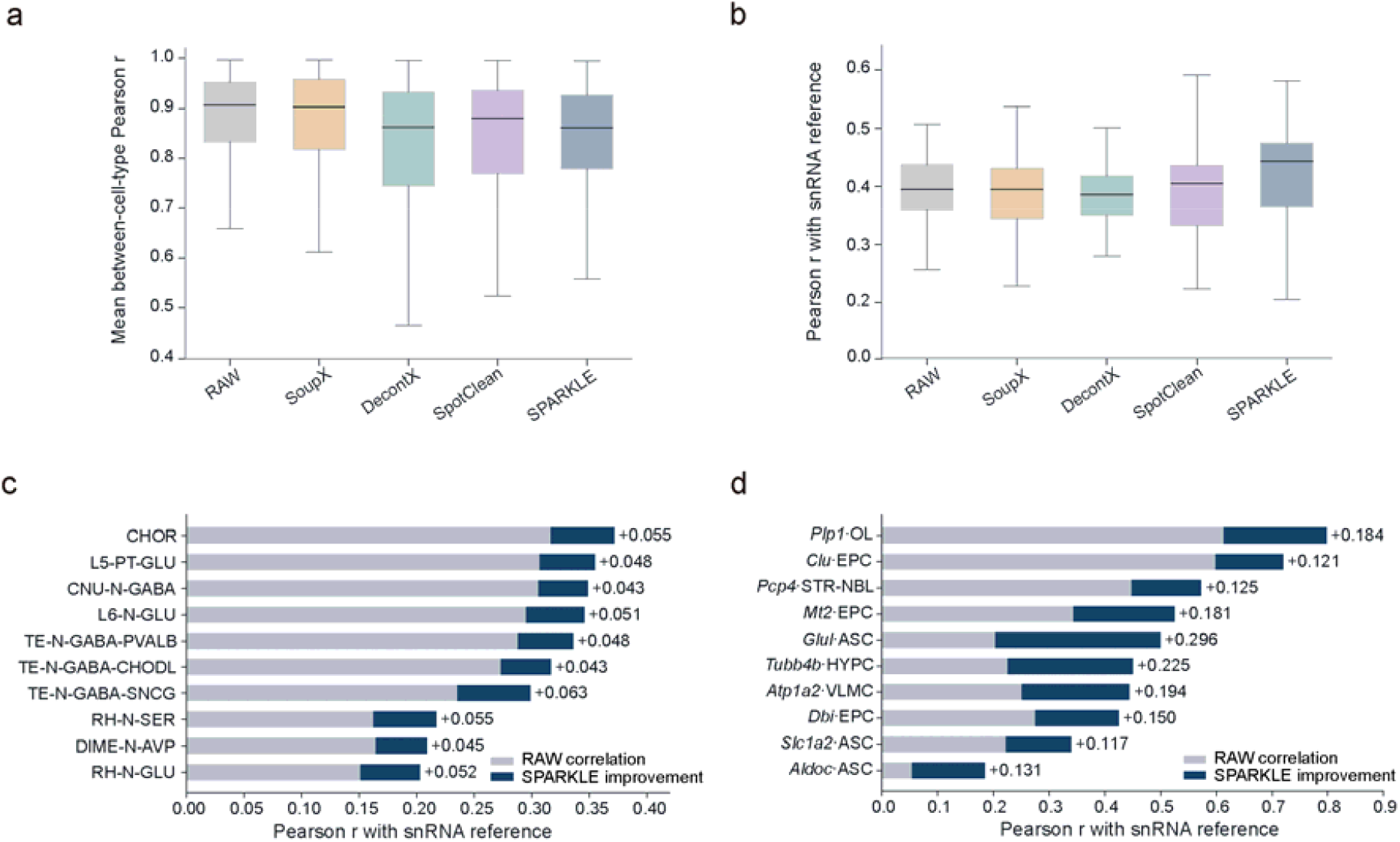
Cell-type separation and concordance with an independent snRNA-seq reference in mouse brain. (a) Pairwise Pearson correlations between cell-type pseudobulks. (b) Correlations between spatial cell-type pseudobulks and matched snRNA-seq reference populations. (c) Top ten cell populations by improvement in reference concordance (SPARKLE versus RAW). (d) Top ten marker-gene/cell-population combinations by improvement in reference concordance.

RCTD [7] with the external reference provided the cell-population assignments used below. Agreement with the external reference improved in 38 of 40 cell populations. The largest gains were TE-N-GABA-SNCG (+0.063), CHOR (+0.055) and RH-N-SER (+0.055) (Fig. 4 c). At the gene level, among marker genes, the largest improvements in global correlation were seen for *Glul* (ASC, +0.296), *Tubb4b* (HYPC, +0.225) and *Atp1a2* (VLMC, +0.194) (Fig. 4 d). SPARKLE therefore reduced shared background between cell types while improving the correspondence of multiple cell populations and marker genes with the independent single-nucleus reference.

### SPARKLE increases tumor-stroma contrast and communication specificity in ovarian cancer

The ovarian-cancer microenvironment contains closely intermingled tumor, stromal and immune cells, providing a stringent setting for testing whether decontamination preserves biological heterogeneity [22, 23]. SPARKLE reduced mean between-cell-type expression correlation from 0.798 in RAW data to 0.664 and increased agreement with the independent scFFPE reference from 0.483 to 0.540, the highest reference agreement among the tested methods; SoupX produced a much lower between-cell-type correlation (0.285) but also reduced reference agreement (0.379), indicating overcorrection rather than improved separation (Fig. 5 a and b).

**Figure 5.**
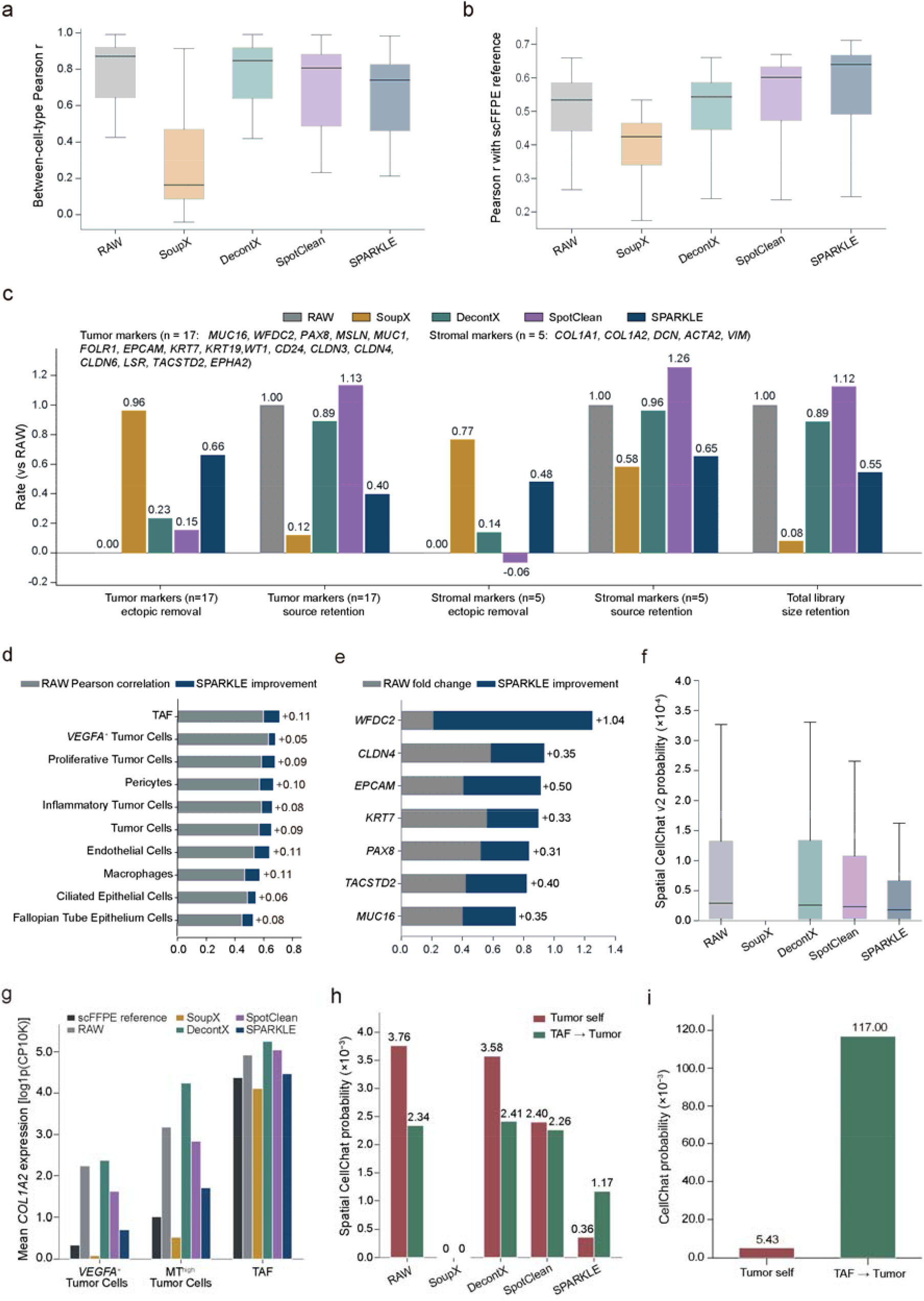
Cell-type contrast, single-cell-reference concordance and *COL1A2-SDC4* communication in ovarian cancer. (a) Pairwise Pearson correlations between cell-type pseudobulks. (b) Correlation between spatial pseudobulks and the independent scFFPE single-cell reference. (c) Ectopic-signal removal rate and source-signal retention in raw counts (versus RAW) for curated tumor and stromal markers, together with total library size retention. (d) Top ten cell populations by improvement in reference concordance (SPARKLE versus RAW). (e) RAW tumor-to-stroma fold differences and SPARKLE increments for representative tumor markers. (f) Spatial CellChat v2 communication probabilities. (g) Mean *COL1A2* expression in *VEGFA*^+^ Tumor Cells, MT^high^ Tumor Cells and TAF. (h) Spatial *COL1A2-SDC4* probabilities for the *VEGFA*^+^ Tumor-cell autocrine and TAF-to-*VEGFA*^+^ paracrine routes. (i) CellChat probabilities for the same routes in the scFFPE reference. TAF, Tumor-Associated Fibroblasts.

Marker-level analysis further revealed changes in tumor-stroma expression contrast. We quantified correction for 17 tumor markers and 5 stromal markers with sufficient leakage evidence as ectopic-signal removal versus source-signal retention in raw counts (Fig. 5 c). SPARKLE removed 66% and 48% of ectopic signal for tumor and stromal markers while retaining 40% and 65% of source expression, respectively. SoupX removed more ectopic signal (96% and 77%) but retained only 12% and 58% of source expression and reduced the total library to 8% of RAW, consistent with overcorrection, whereas DecontX and SpotClean largely preserved source expression but removed less ectopic signal (14-23% for DecontX; 15% of tumor-marker signal and no net removal of stromal-marker signal for SpotClean). Agreement with the scFFPE reference improved in all ten displayed cell populations, with the largest gains of approximately 0.11 in Tumor-Associated Fibroblasts (TAF), endothelial cells and macrophages (Fig. 5 d). Among the tumor markers, *WFDC2* had the largest increase in tumor-to-stroma fold difference (+1.04; Fig. 5 e).

Spatial CellChat communication probabilities were markedly reduced after SPARKLE correction (total probability 0.45 versus 1.02 in RAW data) and were nearly abolished after SoupX correction (0.09), consistent with its removal of over 90% of cellular counts, whereas DecontX (1.03) and SpotClean (0.82) remained close to RAW [8] (Fig. 5 f). This indicated that correction changed ligand-receptor co-expression estimates to different degrees across methods. We examined *COL1A2-SDC4* to determine whether this change also affected the inferred cellular source. In the scFFPE reference, mean *COL1A2* expression was 4.38 in TAF and 0.34 to 1.01 in tumor cells. In spatial RAW data, tumor-cell expression was 2.24 to 3.18, compared with 4.93 in fibroblasts. SPARKLE reduced tumor-cell *COL1A2* to 0.71 to 1.71 while retaining fibroblast expression at 4.47 (Fig. 5 g).

SPARKLE also shifted the dominant inferred *COL1A2-SDC4* communication route from autocrine to paracrine. In RAW data, the autocrine probability for *VEGFA*^+^ Tumor Cells (3.76×10−3) was higher than paracrine signalling from TAF (2.34×10^−3^), and this ordering remained after DecontX (3.58×10^−3^ versus 2.41×10^−3^) and SpotClean (2.40×10^−3^ versus 2.26×10^−3^) correction, whereas neither route was detectable after SoupX correction. After SPARKLE correction, the probabilities were 0.36×10^−3^ and 1.17×10^−3^, respectively, so fibroblast-to-tumor paracrine signalling became dominant. The same ordering was observed in the scFFPE reference (paracrine: 117.00×10^−3^; autocrine: 5.43×10^−3^; Fig. 5 h and i). Because spatial and non-spatial CellChat probabilities are on different scales, we compared the ordering of the two routes rather than their absolute values. In this example, removing fibroblast-derived collagen signal from tumor cells restored a communication source consistent with the single-cell reference.

### Hyperparameter robustness and computational performance

Two SPARKLE hyperparameters determine how out-of-mask spatial information is used: the spatial decay scale λ sets how far a source cell can influence an out-of-mask location, and background-bin size sets the aggregation resolution of out-of-mask observations. We tested whether changing either parameter altered correction performance or the main biological conclusions.

In a simulation with a known true scale, RMSE reduction was 35.2% when λ equalled the true value of 50 µm, which was also selected by the automatic search (Fig. 6 a). RMSE reduction remained between 30.8% and 35.7% for λ values from 30 to 100 µm, but fell to 5.8% at 5 µm and 0.6% at 500 µm. Moderate misspecification therefore had little effect once the main leakage range was covered, whereas severely short or long kernels performed poorly.

**Figure 6.**
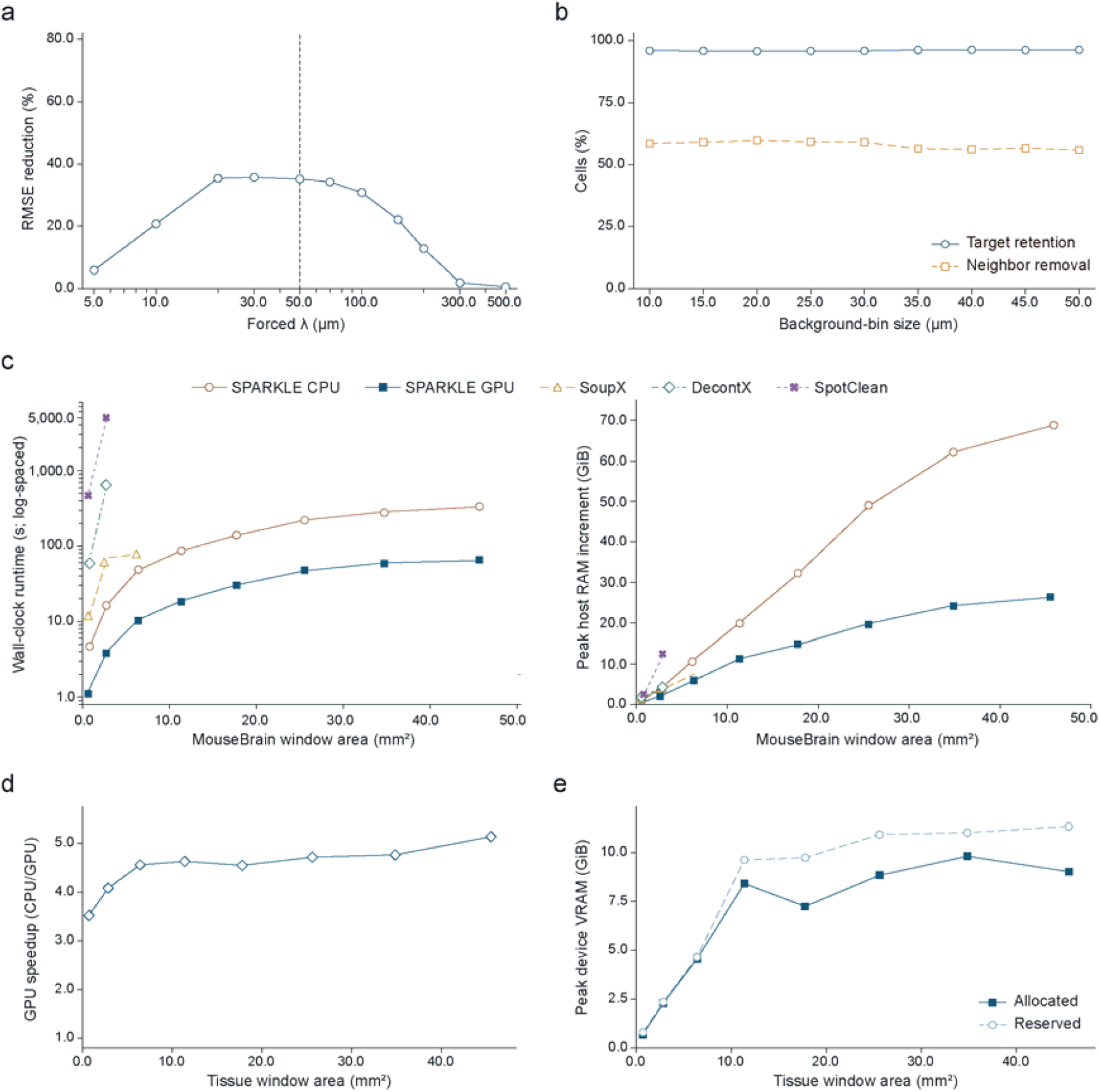
Hyperparameter robustness and computational performance. (a) RMSE reduction across fixed λ values in simulation. (b) Retention of sstIN signal and removal of neighbouring ectopic signal across background-bin sizes. (c) Wall-clock runtime and peak host RAM increment across eight mouse-brain windows of increasing size. (d) GPU speed-up across eight mouse-brain windows of increasing size. (e) Allocated and Reserved GPU memory usage.

Changing the background-bin side length from 10 to 50 µm in the axolotl dataset did not change the main *SST* result. Retention of source-cell signal remained 95.7% to 96.2%, and removal of ectopic signal from neighbouring non-sstIN cells remained 55.9% to 59.7% (Fig. 6 b).

SPARKLE was computationally efficient at both CPU and GPU settings. Across eight mouse-brain windows spanning 0.71 to 45.50 mm^2^ and 1,831 to 71,476 cells, runtime scaled near-linearly with tissue area (power-law exponent 1.08 on CPU and 1.00 on GPU, R2≥0.99). In the largest window, runtime was 330.3 s on CPU and 64.3 s on GPU, while GPU speed-up increased from 3.52-fold to 5.14-fold across window sizes (Fig. 6 c-e). Compared with the baseline methods, SPARKLE was one to three orders of magnitude faster at comparable or lower memory use. On the 2.84 mm^2^ window with 6,804 cells, it completed in 4.1 s on GPU, compared with 69.9 to 5,085.0 s for SoupX, DecontX and SpotClean (Fig. 6 c).

## Discussion

SPARKLE estimates local RNA leakage from out-of-mask counts in the same tissue section and applies the estimate conservatively at cell level. Across ten simulations, SPARKLE improved per-cell expression recovery, attaining the highest cell-wise R^2^ in 8 of 10 scenarios and the lowest RMSE in 9 of 10 scenarios. In axolotl telencephalon, most of the correction occurred around SST source cells while source expression was retained. In mouse brain and ovarian cancer, correction reduced convergence between cell-type profiles and improved agreement with independent single-cell references. The COL1A2-SDC4 example showed that low-level ectopic expression can also change the inferred direction of ligand-receptor communication.

SPARKLE differs from existing approaches in how it uses out-of-mask capture locations. Empty-droplet studies have shown that cell-free observations characterize ambient RNA and reveal its effects on cell identification and expression specificity [15–17, 24]. High-resolution spatial data add coordinates to those observations, making it possible to model background according to nearby source cells and distance rather than composition alone. SPARKLE uses this within-sample spatial variation to constrain the local component eligible for subtraction, complementing methods that address global ambient RNA, cluster-level mixture, spot swapping or reference-guided decomposition [11, 12, 15, 24]. The per-gene goodness-of-fit threshold and protection of highly expressing cells provide two safeguards: when out-of-mask data do not support local leakage for a gene, SPARKLE leaves its original expression unchanged.

We evaluated the real datasets using source-marker retention, ectopic expression in neighbouring cells, cell-type separation and reference concordance. Considering these measures together reduces the chance that a single metric rewards overcorrection or simple sparsification. Independent single-cell references are not absolute ground truth: their profiles depend on dissociation, nuclear versus whole-cell measurements, sequencing depth and cell-state coverage. Reference correlation and marker localization should therefore be interpreted together at the same annotation level. Future benchmarks should prioritize in situ proteins, imaging markers or experimental mixture controls from the same tissue.

The ovarian-cancer *COL1A2-SDC4* analysis gives a concrete example of how local ectopic expression affects mechanistic inference. Fibroblast-derived collagen assigned to Tumor cells can make paracrine signalling appear autocrine. After correction, *COL1A2* remained high in Tumor-associated fibroblasts, decreased in two Tumor-cell populations, and the dominant inferred communication route matched the ordering in the non-spatial scFFPE reference. Communication algorithms and prior resources can change candidate interactions and their ranking [8, 13], so *COL1A2-SDC4* should be treated as a testable hypothesis that requires validation by spatial protein measurements, receptor activation or tissue imaging.

Two model assumptions deserve further work. The shared isotropic λ is interpretable and computationally tractable, but it cannot represent tissue barriers, directional diffusion or region-specific permeabilization. Region-specific kernels constrained by tissue images, cell density or anatomical boundaries could be evaluated by holding out out-of-mask locations or testing cross-region prediction. The second assumption is that the segmentation is correct. Segmentation uncertainty could instead be measured by reporting stability intervals across tools such as Baysor, StarDist and Proseg [6, 27, 28] or by perturbing boundaries. Benchmarks that include directional diffusion, tissue barriers, sources outside the field of view and boundary perturbations would help define when each method is applicable and could extend local leakage estimation from matrix correction to spatial-expression quality control with explicit uncertainty quantification.

This study has several limitations. The simulation generator and the primary model both use an exponential distance kernel, which may favour SPARKLE. Out-of-mask counts are not pure technical negative controls; they may include missed cytoplasm, unsegmented small cells or genuine extracellular RNA. The real-data analyses cover a limited number of tissue windows and need replication across samples, batches and platforms. For samples with low goodness of fit, a selected λ at the search boundary or inadequate out-of-mask coverage, “no correction” should remain a formal result rather than relaxing thresholds after inspection to obtain larger subtractions.

SPARKLE uses out-of-mask observations as within-sample evidence to estimate propagation scale, select genes for correction and protect highly expressing source cells. The method is reference-free, fast and scalable, and it limits subtraction to the local component supported by spatial background data. This provides a bounded approach to expression correction in high-resolution spatial transcriptomics for analyses of cell-type localization, tissue-compartment identification and cell-cell communication.

## Data availability

The axolotl telencephalon Stereo-seq data are available from ARTISTA/STOmicsDB (dataset STDS0000056; project CNP0002068) [18, 19]. Mouse-brain T304 Stereo-seq data and the snRNA-seq reference are available through the Mouse Brain Atlas; the processed dataset is deposited in the Brain Science Data Center (doi:10.12412/BSDC.1699433096.20001), with raw data under accession CNP0003837 [20, 21]. Human ovarian-cancer Visium HD spatial data were obtained from the 10x Genomics Cross-Platform Comparison Visium HD dataset [22]. The independent scFFPE single-cell reference was obtained from the 10x Genomics Human Ovarian Cancer FFPE Single Cell Gene Expression Flex dataset [23]. The simulation generator, analysis-window definitions and processing scripts are provided with the code repository (https://github.com/WangShuai-3/SPARKLE). All third-party data remain subject to their original licences.

## Code availability

SPARKLE is available as the open-source Python package stambient from the Python Package Index (PyPI; https://pypi.org/project/stambient/) and supports Python 3.9 or later. All analyses reported in this study were performed using stambient v0.1.3. Source code, tests, data-preprocessing utilities and scripts used to generate the benchmarks are available from https://github.com/WangShuai-3/SPARKLE. The software is released under the MIT License.

## Author contributions

S.W. conceived the project. X.W. supervised the study and provided overall guidance. S.W. and B.Z. carried out the method development, testing, and data analyses. B.Z. and S.L. performed the visualization. S.W., B.Z. and S.L. wrote the original draft, and X.W. reviewed the manuscript.

## Acknowledgements

We would like to thank DCS Cloud (https://dcs.cloud) for providing the computational resources and software support necessary for this study. We sincerely thank the China National GeneBank (CNGB) for the support they provided.

## Key Points

- SPARKLE uses capture locations outside cell-segmentation masks as within-sample spatial evidence of local RNA leakage.
- A shared propagation scale, gene-specific coefficients and per-gene goodness-of-fit gating restrict correction to the locally supported leakage component.
- Across simulations and three real-tissue datasets, SPARKLE mitigated signal leakage, improved concordance with single-cell references, and enhanced downstream cell-cell communication inference.
- Sensitivity analyses were stable across plausible parameter settings, and resource benchmarks showed near-linear scaling to large tissue windows.

